# Reverse engineering of the pattern recognition receptor FLS2 reveals key design principles of broader recognition spectra against evading flg22 epitopes

**DOI:** 10.1101/2024.10.10.617594

**Authors:** Songyuan Zhang, Songyuan Liu, Hung-Fei Lai, Amedeo Caflisch, Cyril Zipfel

## Abstract

In the ongoing plant-pathogen arms race, plants employ pattern recognition receptors (PRRs) to recognize pathogen-associated molecular patterns (PAMPs), while in successful pathogens, PAMPs can evolve to evade detection. Engineering PRRs to recognize evading PAMPs could potentially generate broad-spectrum and durable disease resistance. In this study, we reverse-engineered two natural FLAGELLIN SENSING 2 (FLS2) variants, VrFLS2XL and GmFLS2b, with extended recognition specificities towards evading flg22 variants. We identified minimal gain-of-function residues enabling blind FLS2s to recognize otherwise evading flg22 variants. We uncovered two strategies: (i) enhancing FLS2-flg22 interaction around flg22’s key evasion sites, and (ii) strengthening direct interaction between FLS2 and its co-receptor BAK1 to overcome weak agonistic and antagonistic flg22s, respectively. Additionally, we leveraged polymorphisms that enhance recognition through unknown mechanisms to engineer superior recognition capability. These findings offer basic design principles for PRRs with broader recognition spectra, paving the way for PRR engineering using precise gene-editing to increase disease resistance in crops.

## Introduction

As the first layer of plant immune recognition, plants employ plasma membrane-localized pattern recognition receptors (PRRs) to detect microbe-/pathogen-associated molecular patterns (MAMPs/PAMPs) of potential invaders and initiate pattern-triggered immunity (PTI). One of the best-studied plant PRRs is the leucine-rich repeat (LRR) receptor kinase FLAGELLIN SENSING 2 (FLS2), which, along with the co-receptor BRASSINOSTEROID INSENSITIVE 1-ASSOCIATED KINASE 1 (BAK1), recognizes flg22 (a 22-amino-acid epitope derived from the N terminus of bacterial flagellin)^1, 2, 3^. FLS2 is ubiquitously present in angiosperms^4^ and plays a crucial role in anti-bacterial disease resistance^5^.

Structural characterization of the FLS2-flg22-BAK1 complex^6^ and previous genetic and biochemical studies^2, 7^ suggest a sequential ligand-induced heterodimerization mechanism of flg22 perception and receptor activation^8^. The flg22 peptide adopts an extended conformation to bind to the concave surface of FLS2’s extracellular leucine-rich repeat domain. Its N-terminal seventeen amino acids (‘address’ segment) interact solely with FLS2, and its C-terminal five amino acids (‘message’ segment) together with FLS2 create a flg22-mediated FLS2-BAK1 interaction interface to recruit BAK1. The FLS2-flg22-BAK1 complex is further stabilized by another direct FLS2-BAK1 interaction interface independent of flg22. The heterodimerization of extracellular domains brings the intracellular kinase domains of FLS2 and BAK1 into proximity enabling trans-phosphorylation and activation of downstream immune signaling^2, 6, 9, 10^, including reactive oxygen species (ROS) production, calcium influx, mitogen-activated protein kinase (MAPK) cascades, transcriptional reprogramming, and callose deposition^11, 12^.

PRR-mediated PAMP recognition imposes strong selective pressures on pathogens, driving them to evade the recognition^13, 14^. For example, many bacterial phytopathogens carry polymorphic flg22 epitopes not recognizable by FLS2^15^. This includes *Agrobacterium tumefaciens*, the causative agent of crown gall disease^3, 16^, *Xanthomonas oryzae* pv. *oryzae* (*Xoo*) and pv. *oryzicola* (*Xoc*), which cause bacterial blight and leaf streak in rice^17^, *X. campestris* pv. *campestris*, responsible for black rot in crucifers^18^, the bacterial wilt disease pathogen *Ralstonia solanacearum*^19^, and *Erwinia amylovora* that causes fire blight^20^. Such immune evasion is also common among the commensal microbiota of healthy plants^21, 22, 23^. Polymorphic flg22 variants apply diverse mechanisms to evade recognition: some act as weak agonists with reduced binding affinity to FLS2, while some act as antagonists that retain interaction with FLS2 but harbor C-terminal mutations to impede BAK1 recruitment. Additionally, another three types of flg22 variants have been reported with atypical evasion behavior through uncharacterized mechanisms^21^.

However, the evolution of non-immunogenic flg22 variants by pathogens can be countered by the co-evolution of FLS2 variants with novel recognition specificities. For example, VrFLS2XL from riverbank grapes (*Vitis riparia, Vr*) can recognize *Agrobacterium tumefaciens* flg22 (flg22^Atum^)^24^, and GmFLS2b of soybean (*Glycine max*, *Gm*) can recognize *Ralstonia solanacearum* flg22 (flg22^Rso^)^25^. Recently, the generation of *Nicotiana benthamiana fls2* mutant has facilitated the rapid characterization of natural FLS2 variants, expanding the known repertoire of FLS2s with novel recognition specificities^26^. For example, QvFLS2 of *Quercus variabilis* (Qv) and TjFLS2 of *Trachelospermum jasminoides* (Tj) are also highly sensitive to flg22^Atum^ when transiently expressed in *N. benthamiana fls2* mutant^26^.

The interfamily transfer of PRRs has proven to be an effective strategy to generate broad-spectrum and potentially durable disease resistance^27, 28, 29^, which holds true not only for phylogenetically-restricted PRRs, like the ELONGATION FACTOR TU RECEPTOR (EFR), but also for the prevalent FLS2. For example, transferring VrFLS2XL to *N. benthamiana* and GmFLS2b to tomato conferred resistance to *A. tumefaciens* and *R. solanacearum*, respectively^24, 25^. However, the broad applicability of PRR transfer is currently limited by the lack of known PRRs, difficulties in identifying new PRRs, the potential for pathogens to evolve evasion, and legal restrictions on the use of transgenic plants.

A promising alternative is the use of precise gene-editing techniques to modify native PRRs, thereby expanding their recognition spectra^30^. The increasingly wider acceptance of genome-edited crops^31^ and the development of CRISPR-mediated precise genome-editing technologies^32^ can enable the fast deployment of engineered PRRs from the lab to the field. However, this approach is currently hindered by our limited knowledge of how to design PRRs with novel recognition specificities. While several previous studies have tried to understand novel specificities of FLS2s, they either did not generate a gain-of-function FLS2^25^ or the resolution of the mapped region was not fine enough to guide precise engineering^20, 24, 33^.

Here, we explored the basic engineering principles behind novel recognition specificities by reverse-engineering two natural FLS2 variants, VrFLS2XL and GmFLS2b. Using a combination of domain swapping, computational design, and DNA shuffling, we identified minimal residues required to enable blind VvFLS2 of cultivated grape (*Vitis vinifera*) and GmFLS2a (an ortholog of GmFLS2b insensitive to flg22^Rso^) to recognize flg22^Atum^ and flg22^Rso^, respectively. Our findings revealed two key engineering principles: (i) enhancing FLS2-flg22 interaction around flg22’s key evading mutations to overcome weak agonistic flg22 variants, and (ii) strengthening direct FLS2-BAK1 interaction to overcome antagonistic flg22 variants. In addition to identifying minimal gain-of-recognition residues, we discovered polymorphic residues outside the flg22- and BAK1-interacting interfaces of GmFLS2b, which influence recognition specificity through unknown mechanisms. By leveraging these residues, we engineered a GmFLS2 variant with enhanced flg22^Rso^ recognition capability. This study advances our understanding on the recognition specificity of FLS2 and reveals fundamental design principles to engineer broader recognition spectra. Additionally, this study provides valuable genetic resources to generate crown gall and bacterial wilt disease resistance in economically important crops.

## Results

### LRR12-19 region of VrFLS2XL is mainly responsible for flg22^Atum^ recognition

The flg22^Atum^ of *A. tumefaciens* was one of the first flg22 variants reported to evade recognition by plants^3^. Previous studies on flg22^Atum^ responsiveness of FLS2 homologs in cultivated grapes (*Vitis vinifera*) and riverbank grapes (*Vitis riparia*) provide an excellent model for investigating how a novel recognition specificity is formed (**Extended Data Fig. 1a**)^16, 24, 34^. Therefore, we aimed to identify minimal key residues from the flg22^Atum^-responsive VrFLS2XL to render the flg22^Atum^- blind VvFLS2 responsive to flg22^Atum^.

We first assessed the flg22^Atum^ responsiveness of VvFLS2 and VrFLS2XL in *N. benthamiana fls2* mutant (**Fig. 1a, b**). As a control, we concurrently tested each FLS2 variant’s response to the canonical flg22 of *Pseudomonas aeruginosa* (flg22^Pa)^ to confirm that the tested FLS2 variant retains its original flg22 recognition capability. Flg22^Atum^ induced ROS production when VrFLS2XL was expressed, but not when VvFLS2 was (**Fig. 1a**). However, VvFLS2 is not completely irresponsive to flg22^Atum^ as weak ROS production could be detected at higher flg22^Atum^ concentrations (**Fig. 1b** and **Extended Data Fig. 1b**). We also tested the flg22^Atum^ responsiveness of other FLS2 homologs, including SlFLS2 from tomato (*Solanum lycopersicum*), AtFLS2 from *Arabidopsis thaliana*, GmFLS2a/b from soybean (*Glycine max*), and NbFLS2 from *N. benthamiana* (**Extended Data Fig. 1c-g**). All these FLS2s were blind to flg22^Atum^, and flg22^Atum^ had no antagonistic effect on flg22^Pa^-induced ROS production. These results categorize flg22^Atum^ as a weak agonist^22^ that evades FLS2 recognition primarily due to loss of binding (consistent with previous binding assays^22, 24, 35^). However, we cannot exclude the possibility that 18Y, 19W, 20S (hereinafter referred to as ^18^YWS^20^) mutations on flg22^Atum^ might also affect BAK1 recruitment, as suggested in previous studies^23, 24^.

**Fig. 1:**
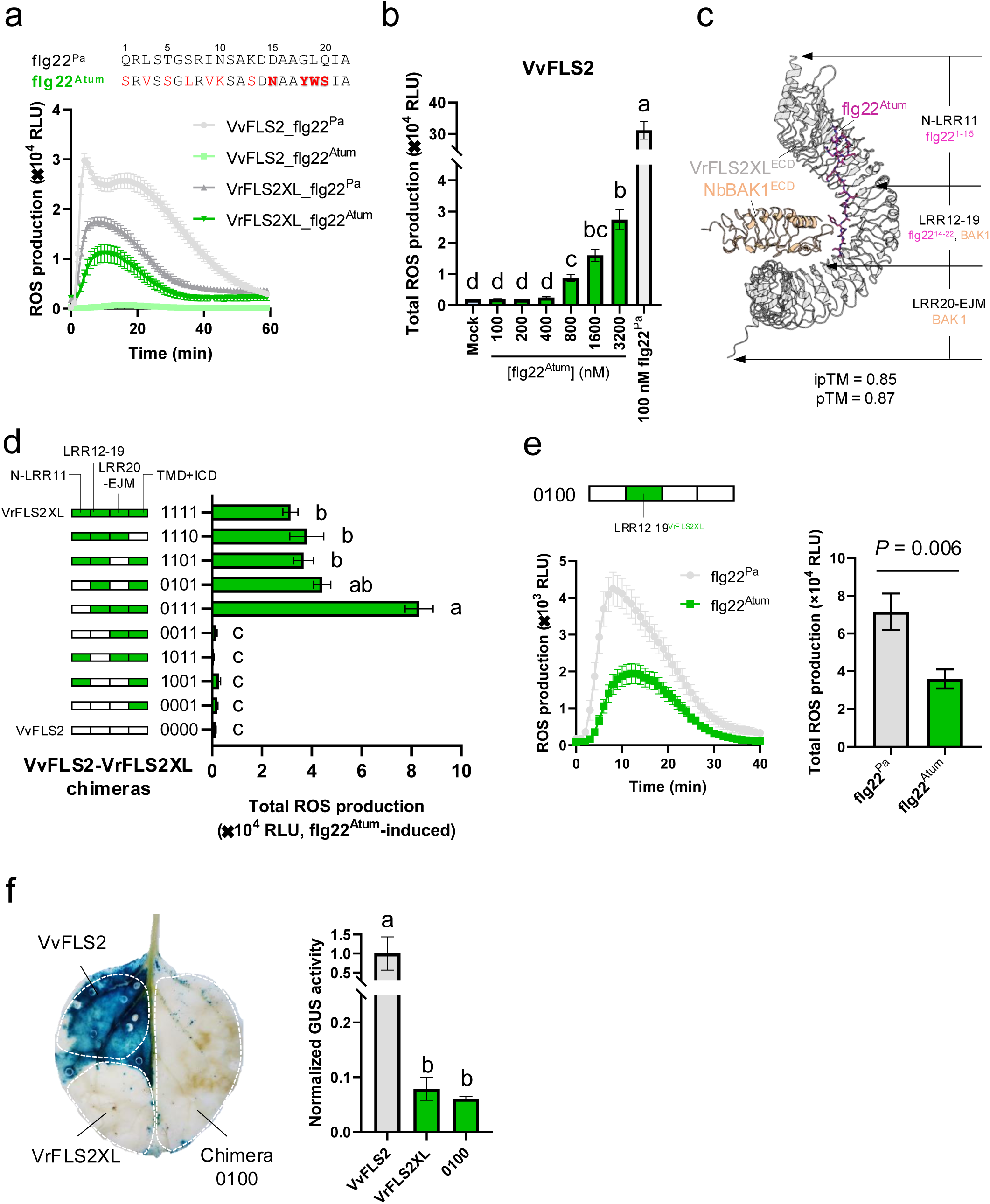
LRR12-19 is mainly responsible for flg22^Atum^ recognition by VrFLS2XL. **a,** Sequences of flg22^Atum^ and flg22^Pa^, and ROS production induced by VvFLS2 and VrFLS2XL in response to these peptides. Mutations in flg22^Atum^ are highlighted in red, with reported key mutations (15N, 18Y, 19W, 20S) bolded and shaded. **b,** Total ROS production triggered by VvFLS2 in response to flg22^Pa^ and increasing concentrations of flg22^Atum^. **c,** AlphaFold3-predicted structure of VrFLS2XL^ECD^-flg22^Atum^-NbBAK1^ECD^ and the segmentation scheme of VrFLS2XL for domain swapping (ECD: extracellular domain; LRR: leucine-rich repeat; EJM: extracellular juxtamembrane domain). Signal peptides were removed from VrFLS2XL and NbBAK1 before uploading sequences for prediction. NbBAK1 is also known as NbSERK3B. **d,** Flg22^Atum^-induced ROS production of VvFLS2-VrFLS2XL chimeras. As an example, ‘0101’ denotes a chimera with its N-LRR11 region (the first digit) and LRR20-EJM region (the third digit) from VvFLS2, while LRR12-19 region (the second digit) and transmembrane domain plus intracellular domain region (TMD+ICD, the fourth digit) from VrFLS2XL. White and green represent VvFLS2 and VrFLS2XL origins, respectively. Signal peptide is included in the N-LRR11 region when constructing chimeric receptors at DNA level, although it is not considered in the structural prediction. **e,** Flg22^Atum^-induced ROS production of chimera ‘0100’. Total ROS production over 40 minutes was calculated. **f,** Histochemical (left) and fluorometric (right) β-glucuronidase (GUS) assays showing the effects of FLS2 variants on restricting the growth of *A. tumefaciens* carrying a GUS reporter gene. Dark blue coloration indicates high GUS activity and a higher degree of *A. tumefaciens* infection. Unless specified, 100 nM of each elicitor was used to trigger responses. Data are shown as mean ± SEM. Kruskal-Wallis test followed by Dunn’s multiple comparisons test was used for statistical analysis in **d** and **f**. Significant differences at *P* < 0.05 are denoted by letters. Mann-Whitney test was used for statistical analysis in **e**.

VvFLS2 and VrFLS2XL exhibit 82.5% protein sequence identity with more than 200 polymorphic residues. To map the minimal sequence determinants for flg22^Atum^ recognition, we first used a chimeric receptor/domain swapping strategy. According to the AlphaFold3-predicted structure of VrFLS2XL^ECD^-flg22^Atum^-NbBAK1^ECD^ complex (**Fig. 1c**) and the solved structure of AtFLS2^ECD^-flg22^Pa^-AtBAK1^ECD^ complex^6^, we divided FLS2 into four fragments: (1) N-LRR11, from the N terminus (signal peptide was included when making DNA constructs, and removed when making the structural prediction) to the 11^th^ LRR, which recognizes the first fifteen amino acids of flg22; (2) LRR12-19, which interacts with amino acids 14-22 of flg22 and provides the first interacting interface for BAK1 recruitment; (3) LRR20-extracellular juxtamembrane (EJM), from LRR20 to the end of the EJM domain, which does not interact with flg22 but includes the direct FLS2-BAK1 interface; and (4) the transmembrane domain and the intracellular domain (TMD+ICD). Using Golden Gate cloning^36^, we constructed a series of chimeric receptors combining fragments of VvFLS2 and VrFLS2XL. As shown in **Fig. 1d** and **Extended Data Fig. 1i**, chimeras responded to flg22^Atum^ in a binary manner. All flg22^Atum^-responsive chimeras carry LRR12-19 of VrFLS2XL, whilst VrFLS2XL with its LRR12-19 replaced by LRR12-19^VvFLS2^ (chimera ‘1011’) could not respond to flg22^Atum^. Conversely, VvFLS2 carrying LRR12-19^VrFLS2XL^ (chimera ‘0100’) gained responsiveness to flg22^Atum^ (**Fig. 1e**). These results indicate that LRR12-19^VrFLS2XL^ is necessary and sufficient for flg22^Atum^ recognition, while other domains might influence the recognition quantitatively.

Interestingly, all flg22^Atum^-responsive chimeras exhibited weaker ROS-induced production to flg22^Pa^ than flg22^Atum^-irresponsive chimeras (**Fig. 1** and **Extended Data Fig. 1h, i**). It is possibly because *A. tumefaciens* remaining on the infiltrated leaves is recognized by flg22^Atum^-responsive chimeric FLS2s, which can reduce the transformation rate as well as deplete signaling components and substrates for ROS production before conducting measurements. This is supported by the observation of occasional weak cell death on young leaves expressing flg22^Atum^-responsive chimeras for more than three days, but not on leaves expressing flg22^Atum^-irresponsive chimeras. It is unlikely that the gain of flg22^Atum^ recognition *per se* affects flg22^Pa^ recognition, as VrFLS2XL has been previously shown to respond to flg22^Pa^ at a similar or stronger level than VrFLS2 when stably expressed in *N. benthamiana* or transiently expressed in *A. thaliana* protoplasts^24^.

Finally, we tested whether chimera ‘0100’ could confer resistance to *A. tumefaciens*. We used *A. tumefaciens* GV3101 carrying a binary plasmid with an intron-containing β-glucuronidase (GUS) gene^37, 38^ to infect *N. benthamiana* leaves transiently expressing VvFLS2, VrFLS2XL, or the chimera ‘0100’. GUS activity served as a proxy for the degree of infection. Both histochemical and fluorometric GUS assays showed that the chimera ‘0100’ effectively restricted *A. tumefaciens* infection to a similar extent as VrFLS2XL (**Fig. 1f**). Collectively, our results show that the LRR12-19 region of VrFLS2XL is primarily responsible for flg22^Atum^ recognition.

### VrFLS2XL residues critical for flg22^Atum^ recognition are associated with key evading mutations of flg22^Atum^

Between VvFLS2 and VrFLS2XL, there are a total of 39 polymorphic sites within the LRR12-19 region (**Fig. 2a**). To identify the minimal key residues for flg22^Atum^ recognition, we first evaluated the roles of polymorphic sites based on the AlphaFold3-predicted structure (**Fig. 1c** and **Fig. 2b**). Given that no polymorphic site was predicted to directly interact with BAK1, we grouped the 39 polymorphic sites into four categories based on their predicted distance from flg22^Atum^ (**Fig. 2c**): (1) residues directly interacting with flg22^Atum^ at a distance less than 4 Å, (2) residues putatively in contact with flg22^Atum^ potentially through long-distance (4-8 Å) electrostatic interactions and hydrogen bond network (if considering the existence of water molecules), (3) neighboring residues of the categories 1 and 2 residues potentially stabilizing the local conformation (although these neighboring residues may not directly contribute to binding interactions, they appear to support the ligand binding by stabilizing the geometric conformation of local motifs, such as the distance between repeat modules and protein curvatures^39, 40^), and (4) residues outside the predicted interaction interface (distance > 8 Å).

**Fig. 2:**
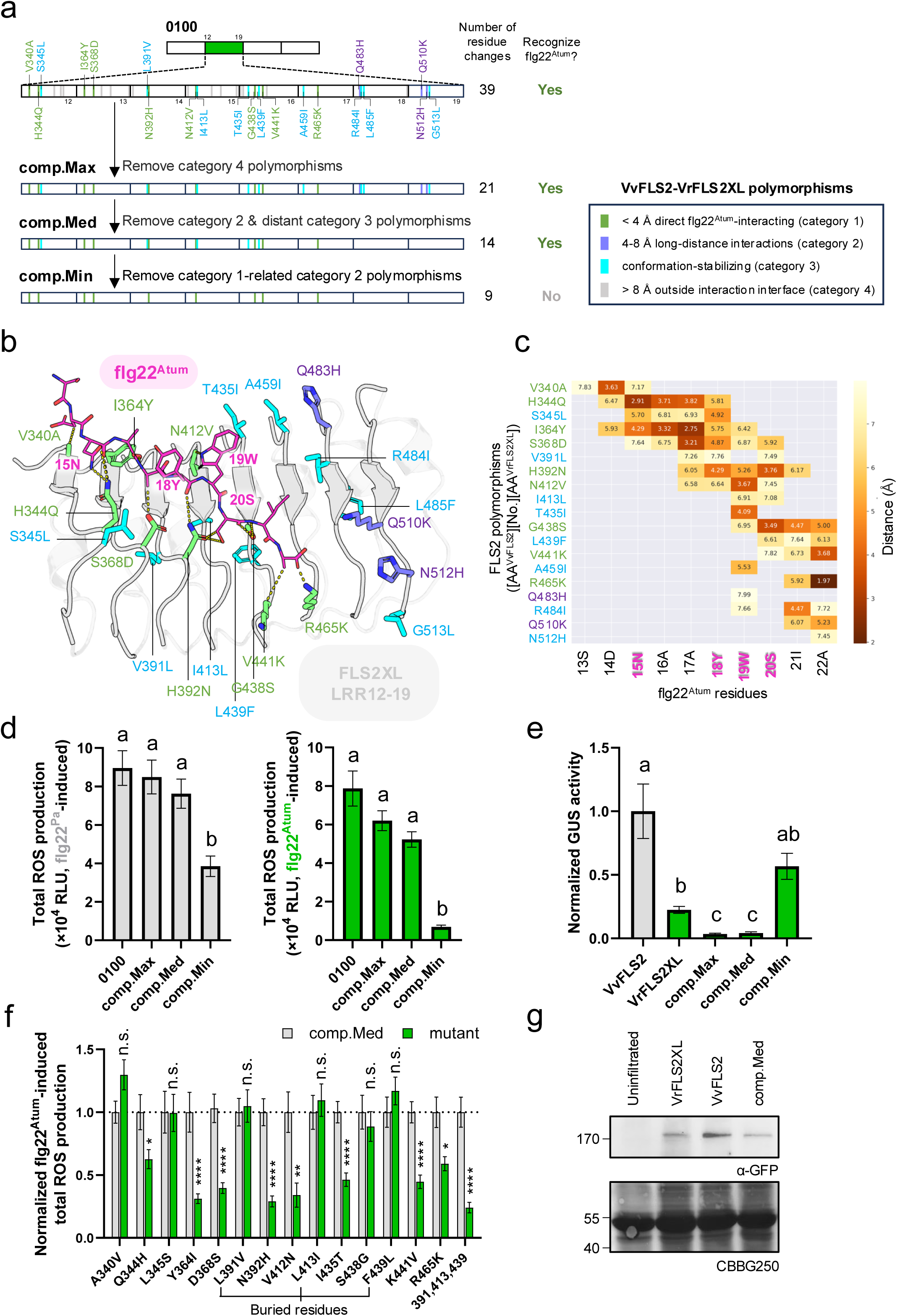
Computer-aided identification and characterization of minimal key residues of VrFLS2XL responsible for flg22^Atum^ recognition. **a,** Schematic overview of polymorphic sites within LRR12-19^VrFLS2XL^ in comparison to VvFLS2, and the process to identify minimal key residues. To label a polymorphic site, the first letter indicates residue status in VvFLS2 and the second letter for VrFLS2XL. The number indicates the residue position in VrFLS2XL. Labels and legends are shared by panel **a-c**. **b,** AlphaFold3-predicted interaction interface of LRR12-19^VrFLS2XL^ and flg22^Atum^. Potential flg22^Atum^-interacting polymorphic residues are displayed with side chains and color-coded. **c,** Contact map showing all interaction pairs with distances < 8 Å between flg22^Atum^ residues and VrFLS2XL polymorphic sites. Numbers indicate the distances. AA: amino acid. **d,** Comparison of total flg22^Pa^- and flg22^Atum^- induced ROS production of VvFLS2 variants listed in panel **a**. **e,** Results of fluorometric GUS assay showing restriction of *A. tumefaciens* infection by natural and engineered FLS2 variants. **f,** Characterization of individual residues in ‘comp.Med’ by mutating each of them from VrFLS2XL to VvFLS2. Total flg22^Atum^-induced ROS production was normalized by the average value of ‘comp.Med’. ‘391,413,439’ denotes three mutations L391V, L413I, and F439L at buried sites. The dotted line indicates Y = 1. **g,** Protein accumulation levels of GFP-tagged VvFLS2, VrFLS2XL, and ‘comp.Med’. Unless specified, 100 nM of each elicitor was used to trigger responses. Total ROS production over 40 minutes was calculated. Data are shown as mean ± SEM. Kruskal-Wallis test followed by Dunn’s multiple comparisons test was used for statistical analysis in **d** and **e**. Significant differences at *P* < 0.05 are denoted by letters. Mann-Whitney test was used for statistical analysis in **f**. The ‘n.s.’ denotes no significant difference. One to four asterisks represent *P* < 0.05, 0.01, 0.001, and 0.0001, respectively.

We then refined the polymorphic sites in a stepwise manner by mutating VrFLS2XL residues to VvFLS2 residues (**Fig. 2a, d**). First, we omitted polymorphic residues outside the interaction interface (Category 4), generating a variant called ‘comp.Max’. ‘Comp.Max’ exhibited only a slight and insignificant reduction in flg22^Atum^ and flg22^Pa^ recognition capabilities compared to chimera ‘0100’. Then, we created ‘comp.Med’ by omitting Category 2 residues and distant Category 3 residues from ‘comp.Max’. This step again resulted in a minimal decrease in both flg22^Pa^ and flg22^Atum^ recognition capabilities. However, when we further omitted neighboring polymorphic residues of Category 1 residues to generate ‘comp.Min’, we observed a significant reduction in both flg22^Pa^ and flg22^Atum^ recognition capabilities (**Fig. 2d** and **Extended Data Fig. 2a**). In terms of conferring resistance to *A. tumefaciens*, ‘Comp.Max’ and ‘comp.Med’ are comparable and even superior to VrFLS2XL, while ‘comp.Min’ exhibited very weak resistance with no significant difference from VvFLS2 **(Fig. 2e)**. The gain of flg22^Atum^ recognition by ‘comp.Med’ is not caused by an increased protein accumulation level (**Fig. 2g**).

To further refine ‘comp.Med’, we first evaluated each LRR carrying polymorphic residues (**Extended Data Fig. 2b**). Given that every LRR of LRR12-17 seemed to be indispensable for flg22^Atum^ recognition, we then individually characterized each polymorphic residue of ‘comp.Med’ (**Fig. 2f** and **Extended Data Fig. 2c**). Most solvent-exposed residues are important for flg22^Atum^ recognition, except 340A and 345L. N392H and K441V mutations affect not only flg22^Atum^ recognition but also flg22^Pa^ recognition (**Extended Data Fig. 2c**). For buried residues (391L, 413L, and 439F), while single mutation did not affect flg22^Atum^ recognition, mutating all of them abolished flg22^Atum^ recognition and compromised flg22^Pa^ recognition. Interestingly, several key residues found in VrFLS2XL are also present in other reported flg22^Atum^-responsive FLS2 variants^26^ (**Extended Data Fig. 2d**). For example, 364Y and 438S are found in all five reported flg22^Atum^- responsive FLS2 variants^26^, while 392N and 412V appear in three of them, implying that plants might have independently evolved similar mechanisms to recognize flg22^Atum^.

Notably, all polymorphic residues critical for flg22^Atum^ recognition are predicted to be located within the flg22^Atum^-interacting interface (**Fig. 2b, c**). Most of them are predicted to directly interact with key evading mutations of flg22^Atum^ (15N, ^18^YWS^20^) or adjacent residues. The exceptions are 441K and 465K, which are predicted to interact with the C-terminus of flg22^Atum^, likely through electrostatic interactions, which may further strengthen the binding of flg22 to FLS2. These findings suggest that enhancing FLS2-flg22 interaction around flg22’s key evasion sites could be a viable strategy for engineering recognition towards weak agonistic flg22 variants.

### LRR20-EJM region of GmFLS2b is mainly responsible for the recognition of antagonistic flg22^Rso^ through modulating direct interaction with BAK1

Instead of losing binding to FLS2, some flg22 variants retain the ability to interact with FLS2 but impede BAK1 recruitment, acting as antagonists of FLS2-induced signaling. A notable example is flg22^Rso^, which was reported to be a weak antagonist for AtFLS2 and a strong antagonist for SlFLS2^21, 33^. The ^18^AYA^20^ motif and 21A mutations have been identified as critical for this antagonism (**Figure 3a**)^25, 33^.

**Fig. 3:**
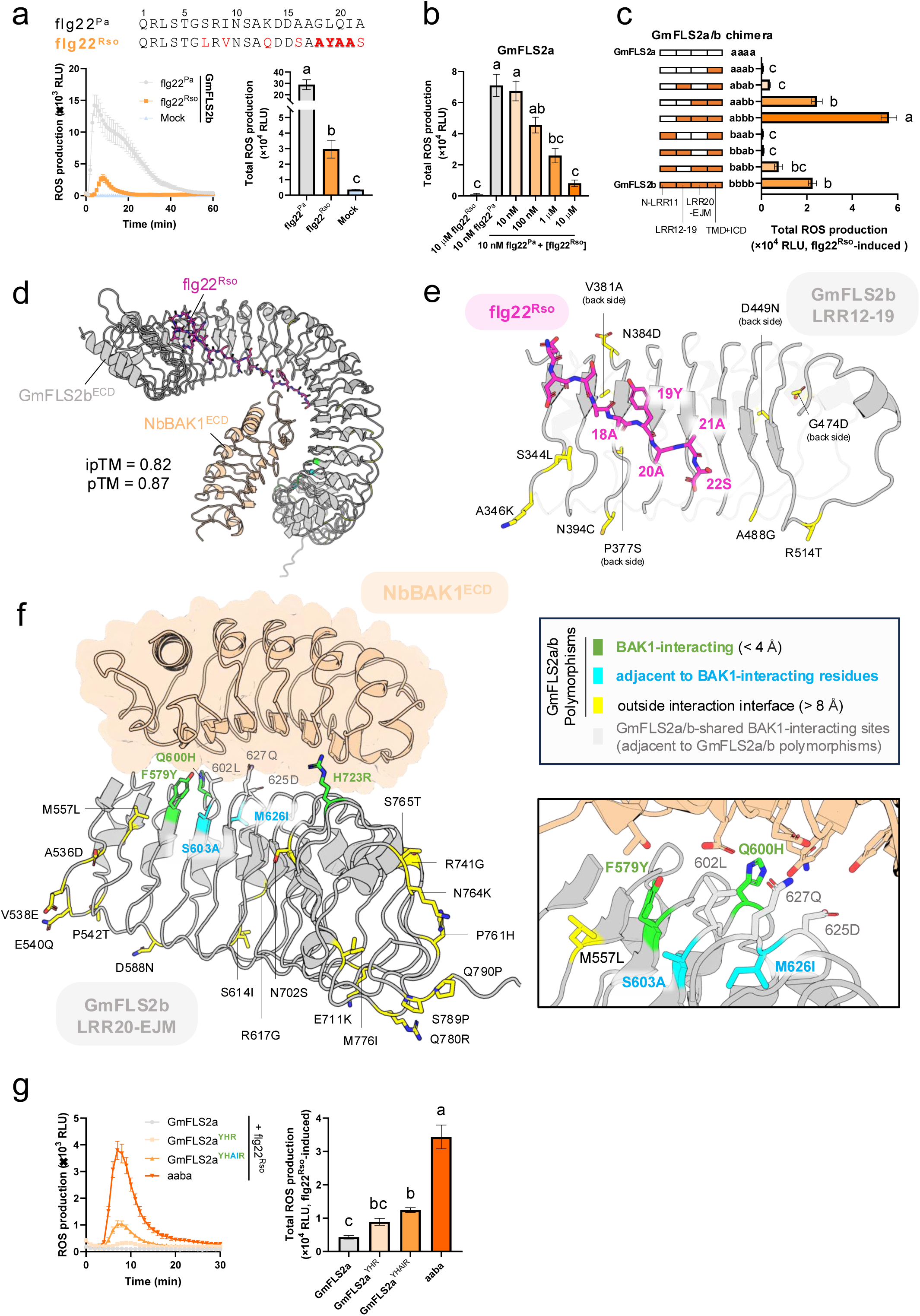
Domain swapping and structural modeling reveal critical polymorphic residues on the direct FLS2-BAK1 interaction interface of LRR20-EJM^GmFLS2b^ for antagonistic flg22^Rso^ recognition. **a,** Sequences of flg22^Rso^ and flg22^Pa^, and GmFLS2b’s ROS responses to them. Mutations in flg22^Rso^ are marked in red. Reported key evading mutations (^18^AYA^20^ and 21A) are in bold and shaded. **b,** ROS responses of GmFLS2a induced by flg22^Rso^ and flg22^Pa^ with increasing concentrations of flg22^Rso^. In the antagonism assay, flg22^Pa^ and flg22^Rso^ were co-administered at the same time. **c,** Flg22^Rso^-induced ROS responses of GmFLS2a/b chimeras. As an example, ‘abab’ denotes a chimera with its N-terminus to the 11^th^ leucine-rich repeat (N-LRR11, the first letter) region and 20^th^ LRR to extracellular juxtamembrane domain (LRR20-EJM, the third letter) region from GmFLS2a, while LRR12-19 region (the second letter) and transmembrane domain plus intracellular domain (TMD+ICD, the fourth letter) from GmFLS2b. White and orange represent GmFLS2a and GmFLS2b origins, respectively. **d,** Overview of the AlphaFold3 (AF3)-predicted GmFLS2b^ECD^-flg22^Rso^-NbBAK1^ECD^ complex structure. Signal peptides were removed from GmFLS2b and NbBAK1 before uploading sequences for prediction. ECD: extracellular domain. **e,** AF3-predicted structure of GmFLS2b^LRR12-19^ in complex with C-terminal flg22^Rso^. All polymorphic sites lie outside the interaction interface with flg22 and BAK1. **f,** AF3-predicted structure of GmFLS2b^LRR20-EJM^-NbBAK1^ECD^ complex. The left panel provides an overview; the right panel zooms in on the direct FLS2-BAK1 interaction interface. Polymorphic residues are displayed with side chains and color-coded in green, cyan, and yellow based on categories from the legends. Three direct BAK1-interacting residues with no difference between GmFLS2a/b are also shown because they are adjacent to polymorphic sites. **g,** Flg22^Rso^-induced ROS responses of GmFLS2a carrying predicted critical residues, compared to GmFLS2a and chimera ‘aaba’. Total ROS production over 30 minutes was calculated. Abbreviations: YHR (F579Y, Q600H, H723R) and YHAIR (F579Y, Q600H, S603A, M626I, H723R). Unless specified, 100 nM of each elicitor was used to trigger responses, with total ROS measured as relative light units (RLU) accumulated over 40 minutes. Data are shown as mean ± SEM. Kruskal-Wallis test followed by Dunn’s multiple comparisons test was used for statistical analysis in **a**, **b**, **c**, and **g**. Significant differences at *P* < 0.05 are denoted by letters.

GmFLS2b was the first FLS2 variant reported to recognize flg22^Rso^^25^. In contrast, GmFLS2a, another copy of soybean FLS2 with 90.6 % protein sequence identity to GmFLS2b, does not respond to flg22^Rso^ when expressed alone in *N. benthamiana*. Before studying them, we first validated the flg22^Rso^ responsiveness of GmFLS2a and GmFLS2b using the *N. benthamiana fls2* mutant. GmFLS2b could recognize flg22^Rso^, but the ROS production induced by flg22^Rso^ was significantly weaker than that induced by flg22^Pa^ (**Fig. 3a**). GmFLS2a, however, was irresponsive to flg22^Rso^, even at a high concentration (10 μM), and flg22^Rso^ acted as an antagonist of flg22^Pa^-induced ROS production for GmFLS2a (**Fig. 3b** and **Extended Data Fig. 3a**).

To map the minimal sequence determinants required for flg22^Rso^ recognition by GmFLS2b, we first performed domain swapping between GmFLS2a and GmFLS2b, using the same segmentation scheme as in the VrFLS2XL case. The resulting chimeras exhibited varying levels of flg22^Rso^ recognition (**Fig. 3c** and **Extended Data Fig. 3b**); though they responded similarly to flg22^Pa^ (**Extended Data Fig. 3b, c**). Notably, the chimera ‘aabb’, which contains the LRR20-EJM domain of GmFLS2b, responded to flg22^Rso^ at a level comparable to GmFLS2b, indicating that LRR20-EJM^GmFLS2b^ is mainly responsible for flg22^Rso^ recognition. While LRR12-19^GmFLS2b^ only rendered a slight and statistically insignificant ROS response to flg22^Rso^ (chimera ‘abab’), its combination with LRR20-EJM^GmFLS2b^ (chimera ‘abbb’) significantly enhanced flg22^Rso^ recognition. Interestingly, N-LRR11^GmFLS2b^ negatively influenced flg22^Rso^ perception, as chimeras lacking N-LRR11^GmFLS2b^ exhibited a stronger flg22^Rso^-induced ROS production than those containing it. The TMD and ICD did not appear to affect flg22^Rso^ perception (**Extended Data Fig. 3d**).

Since the LRR20-EJM region interacts exclusively with BAK1 and antagonistic flg22^Rso^ impedes BAK1 recruitment, we hypothesized that specific BAK1-interacting residues within LRR20-EJM^GmFLS2b^ could strengthen the FLS2-BAK1 interaction to overcome flg22^Rso^ antagonism. Based on the AlphaFold3-predicted structure (**Fig. 3d, f**), three polymorphic residues—F579Y, Q600H, and H723R—directly interact with NbBAK1. However, these changes were insufficient to generate statistically significant ROS production in response to flg22^Rso^ (GmFLS2a^YHR^ in **Fig. 3g**). Building on insights from the VrFLS2XL case, we introduced two additional mutations, S603A and M626I, adjacent to the direct interaction sites shared by GmFLS2a/b (602L, 627Q, and 625D). This resulted in weak but statistically significant ROS production in response to flg22^Rso^ (GmFLS2a^YHAIR^ in **Fig. 3g**).

Given LRR12-19^GmFLS2b^’s significant contribution to flg22^Rso^ recognition, we also examined the predicted structure of LRR12-19^GmFLS2^ (**Fig. 3e**). Surprisingly, all polymorphic sites within LRR12-19^GmFLS2b^ are predicted to be outside the flg22- or BAK1-interacting interface, suggesting that these residues may influence recognition specificity through unknown mechanisms, for example, modulating interactions with other regulators.

### DNA shuffling identifies minimal gain-of-recognition residues and generates greater recognition capacity towards flg22^Rso^

The observation that GmFLS2a^YHAIR^ did not respond to flg22^Rso^ as strongly as the chimera ‘aaba’ (**Fig. 3g**) suggests the presence of additional polymorphic residues contributing to flg22^Rso^ recognition. To identify such residues and further narrow down the minimal gain-of-recognition residues, we employed DNA shuffling^41^, a method that has been successfully used to study plant immune receptors like Cf-4, Cf-9, and Pto^42, 43^. As shown in **Fig. 4a**, DNA shuffling could generate numerous variants with different combinations of polymorphic residues from LRR20-EJM^GmFLS2a/b^. By characterizing these variants, we would be able to determine the contribution of each polymorphic residue to flg22^Rso^ recognition. To better use it in our case, we optimized the classic DNA shuffling method in two ways. First, we made it compatible with Golden Gate cloning, allowing seamless integration with the chimeric receptor construction strategy. Second, we employed the broad-host-range negative selection marker *SacB*^44^ to enable direct library construction in *A. tumefaciens* and meanwhile reduced the occurrence of false-positive clones in the resulting library. Based on the characterization of thirty variants generated by DNA shuffling (**Supplementary Table 3**), we analyzed how each polymorphic site contributes to flg22^Rso^ recognition (**Fig. 4b**). The results of DNA shuffling indicate that 600H and 603A of GmFLS2b are indispensable for flg22^Rso^ recognition. To assess the sufficiency and necessity of these two residues, we introduced Q600H and S603A mutations individually or together to GmFLS2a (**Fig. 4c** and **Extended Data Fig. 4a**). S603A alone did not enable GmFLS2a to respond to flg22^Rso^. While Q600H could sometimes confer a weak response to flg22^Rso^, it was not consistently statistically significant across replicates or in different contexts (e.g., GmFLS2a^YHR^ in **Figure 3f**). However, the combination of Q600H and S603A generated stable ROS production in response to flg22^Rso^, which was not significantly weaker than that of GmFLS2b (**Fig. 4e** and **Extended Data Fig. 4e**). The gain of flg22^Rso^ recognition was not due to a change in FLS2 abundance (**Fig. 4f**) and was not accompanied by an increase in flg22^Pa^ responsiveness (**Extended Data Fig. 4b, d, f**). We further validated these findings by introducing H600Q and A603S mutations into the chimera ‘aaba’ (**Fig. 4d** and **Extended Data Fig. 4c**). H600Q completely abolished flg22-induced ROS production, while A603S significantly reduced it. It has been previously reported that some flg22 variants can uncouple immune outputs; for example, triggering only a reduced ROS response but no other immune responses^21^. Therefore, we assessed calcium influx and MAP kinase activation mediated by GmFLS2a^H600Q, S603A^. Both immune outputs can be induced by flg22^Rso^ (**Fig. 4g, h**). Collectively, these results demonstrate that H600Q and S603A are *bona fide* minimal gain-of-function mutations for GmFLS2a to recognize flg22^Rso^.

**Fig. 4:**
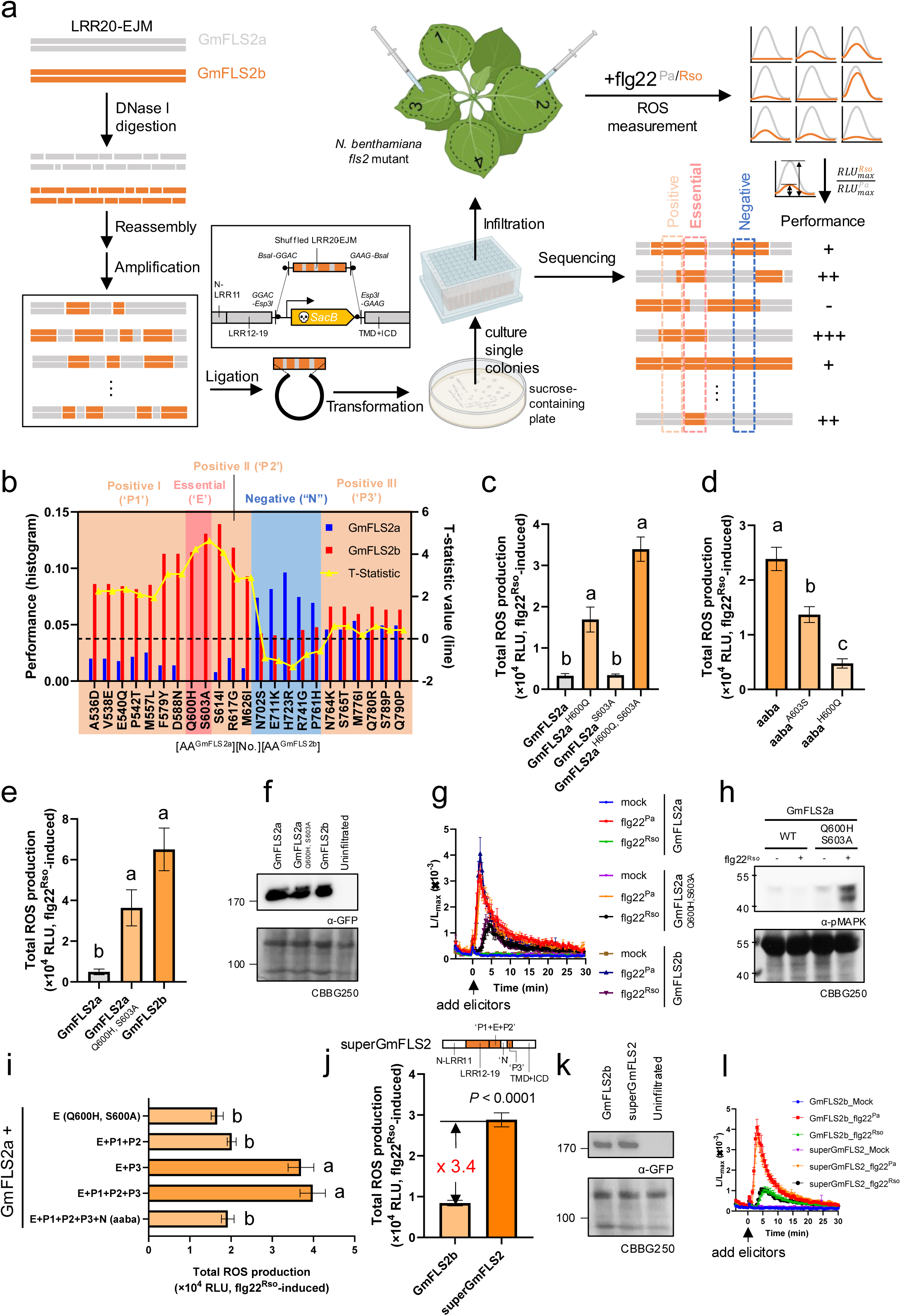
DNA shuffling dissects the roles of polymorphisms within the LRR20-EJM region of GmFLS2a/b and generates greater recognition capacity towards flg22^Rso^. **a,** Schematic diagram of the DNA shuffling workflow. LRR20-EJM^GmFLS2a^ and codon-optimized LRR20-EJM^GmFLS2b^ (to increase its homology with GmFLS2a) were used as parental templates. After standard DNA shuffling procedures, shuffled LRR20-EJM variants were introduced to a vector containing the other three segments of GmFLS2a and *SacB* counterselection marker, via Golden Gate cloning. The Golden Gate product was purified and electroporated into *A. tumefaciens*. Transformed cells were plated on LB plates containing sucrose. Single colonies were cultured in 96-well plates and infiltrated to *N. benthamiana fls2* mutant. Meanwhile, colonies were sequenced. Three days after infiltration, flg22^Pa^- and flg22^Rso^-induced ROS responses of each shuffled variant were measured. The performance of each variant was quantitatively evaluated by the ratio of maximal RLU values of flg22^Pa^ and flg22^Rso^-induced ROS bursts. Variants showing no visible flg22^Rso^-induced ROS burst were assigned a performance of zero. Analysis of the correlation between genotype and phenotype could identify polymorphic residues with different functions. **b,** Analysis of DNA shuffling results. See the Methods section for detailed explanations of this plot. In brief, a positive/negative T-statistic value indicates that the GmFLS2b residue at this site potentially makes a positive/negative contribution to flg22^Rso^ recognition. The highest T-statistic values at Q600H and S603A sites suggest that 600H and 603A are essential. The H723R site has the lowest T-statistic value, implying a likely negative effect on flg22^Rso^ recognition. Polymorphic sites are divided into five groups based on their location and predicted effects on flg22^Rso^ recognition: Groups ‘P1’, ‘P2’, and ‘P3’ potentially act as enhancers. Group ‘N’ potentially acts as a suppressor. Group ‘E’ is indispensable. **c,** Flg22^Rso^-induced ROS production of GmFLS2a with single and double mutations of H600Q and S603A. **d,** Effects of A603S and H600Q on the flg22^Rso^-induced ROS response of chimera ‘aaba’. **e-g,** Comparisons of Flg22^Rso^-induced ROS production (**e**), protein abundance (**f**), and flg22^Pa^/flg22^Rso^-induced calcium influx (**g**) between GmFLS2a, GmFLS2a^H600H, S603A^, and GmFLS2b. **h,** Flg22^Rso^-induced mitogen-activated protein kinase (MAPK) phosphorylation of GmFLS2a and GmFLS2a^Q600H, S603A^. Flg22^Rso^ was syringe-infiltrated into intact leaves, and samples were taken 15 minutes after infiltration. **i,** Effects of DNA shuffling-predicted ‘enhancer’ and ‘suppressor’ polymorphisms on flg22^Rso^-induced ROS production. See panel **b** for the definitions of ‘E’, ‘P1’, ‘P2’, ‘P3’, and ‘N’. **j-l,** Comparisons of flg22^Rso^-induced total ROS production (**j**), protein accumulation (**k**), and flg22^Pa^/flg22^Rso^-induced calcium influx (**l**) between GmFLS2b and ‘superGmFLS2’. ‘SuperGmFLS2’ combines all the GmFLS2b-originated (orange) regions/polymorphic residues with positive effects on flg22^Rso^ recognition. Unless otherwise specified, 100 nM of elicitor was used to trigger ROS responses, and total ROS production represents the sum of relative light units (RLU) over 40 minutes. Data are presented as mean ± SEM. Kruskal-Wallis test followed by Dunn’s multiple comparisons test was used for statistical analysis in **c**, **d**, **e**, and **i**, with significant differences at *P* < 0.05 indicated by letters. Mann-Whitney test was used for **j**.

In addition to the two essential residues, the DNA shuffling results suggest that other polymorphic residues function as suppressors or enhancers to affect flg22^Rso^ recognition. According to their predicted functions, we categorized them into the following groups: Positive I (‘P1’, from N536D to N588D), Positive II (‘P2’, from S614I to M626I), Negative (‘N’, from N702S to P761H), and Positive III (‘P3’, from N764K to Q790P) (**Fig. 4b**). To verify the DNA shuffling-predicted effects of each group, we sequentially introduced them into GmFLS2a^Q600H, S603A^ (GmFLS2a with group ‘E’) (**Fig. 4i** and **Extended Data Fig. 4g**). The introduction of ‘P1’ and ‘P2’ resulted in a slight, but not significant, enhancement of ROS production in response to flg22^Rso^, while ‘P3’ had a significant positive effect. ‘P3’ also significantly enhanced flg22^Pa^ responsiveness (**Extended Data Fig. 4h**); though, not as markedly as to flg22^Rso^. The subsequent introduction of ‘N’ significantly decreased flg22^Rso^-induced but not flg22^Pa^-induced ROS production. Further analysis revealed that the BAK1-interacting polymorphic residue H723R is primarily responsible for the suppressive effect of the ‘N’ region, as introducing the R723H mutation into the chimera ‘aaba’ increased ROS production in response to flg22^Rso^ (**Extended Data Figure 4i-k**). However, the specific mechanisms by which ‘P3’ polymorphic residues influence flg22^Rso^ recognition remain unclear, as they are located in the C-terminal capping domain of the LRRs and the extracellular juxtamembrane domain (**Fig. 3b**) – regions that have not been previously studied. These findings underscore the significant role of ‘dark matter’ polymorphisms in modulating recognition specificity. The consistency between DNA shuffling predictions and subsequent experimental results highlights DNA shuffling as a powerful tool for exploring novel recognition specificities of PRRs.

By excluding the two identified suppressive regions and combining all ‘enhancer’ regions, we engineered a variant termed ‘superGmFLS2’, which exhibited a more than 3-fold increase in total ROS production in response to flg22^Rso^ compared to wild-type GmFLS2b (**Fig. 4j** and **Extended Data Fig. 4l**). There was also a slight enhancement in ROS production in response to flg22^Pa^ (**Extended Data Fig. 4m**), perhaps due to the ‘P3’ region discussed before (**Extended Data Fig. 4h**). The enhanced ROS production is not because of an increased FLS2 expression level (**Figure 4k**). However, we did not detect a significant increase in calcium influx (**Figure 4l**). Whether ‘superGmFLS2’ can confer stronger resistance to *R. solanacearum* than GmFLS2b remains to be tested in the future.

### The broad applicability of FLS2 engineering principles relies on the evasion mechanisms employed by flg22 variants

We then investigated whether the insights gained from reverse-engineering VrFLS2XL and GmFLS2b could be broadly applied to engineer FLS2s from other plants. Leveraging the modularity of our Golden Gate-based chimeric receptor construction strategy, we efficiently transplanted the LRR12-19 region from VrFLS2XL and the LRR20-EJM region from GmFLS2b into various FLS2 homologs irresponsive to flg22^Atum^ and flg22^Rso^ (**Extended Data Fig. 1c-g** and **Extended Data Fig. 5**), replacing their native counterparts.

Transplanting LRR12-19^VrFLS2XL^ enabled AtFLS2, SlFLS2, GmFLS2a, and GmFLS2b to gain recognition to flg22^Atum^. However, NbFLS2 lost its ability to detect canonical flg22^Pa^ following this transplantation (**Fig. 5a**). Additionally, the ROS production kinetics induced by AtFLS2 with transplanted LRR12-19^VrFLS2XL^ exhibited unusual patterns, possibly due to incompatibilities between residues from the two different origins at the junctions. Interestingly, GmFLS2b with LRR12-19^VrFLS2XL^ maintained a reduced capability to recognize flg22^Rso^ while gaining recognition of flg22^Atum^. These findings suggest a generalizable design principle: engineering FLS2 recognition towards weak agonistic flg22 variants can be achieved by enhancing FLS2-flg22 interactions around key evasion sites (**Fig. 5c, d**).

**Fig. 5:**
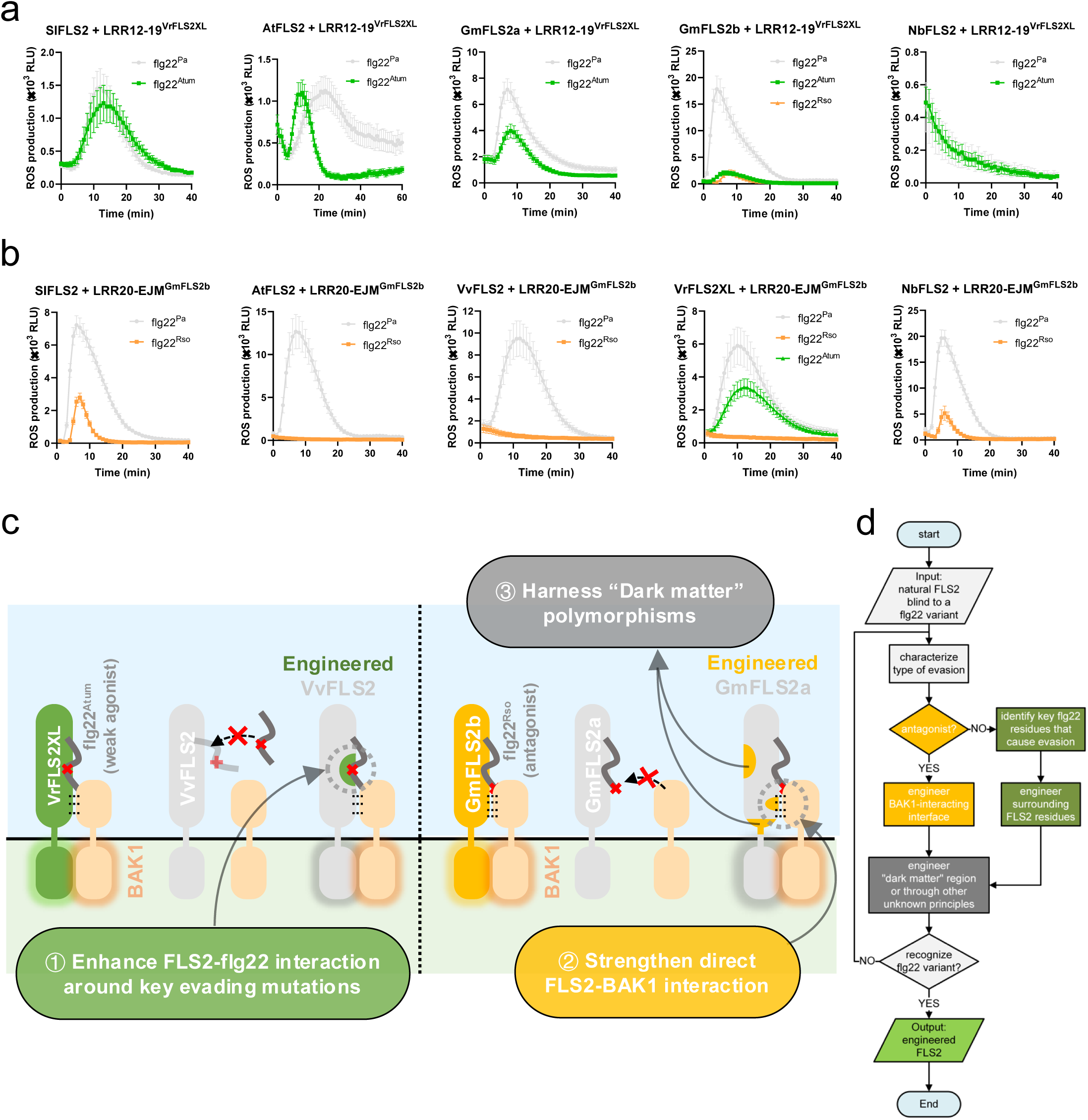
Summary of FLS2 engineering principles and their universal applicability. **a,** ROS responses of FLS2 homologs after transplantation of LRR12-19^VrFLS2XL^. **b,** ROS responses of FLS2 homologs after transplantation of LRR20-EJM. **c,** Summary of proposed design principles to engineer expanded recognition spectra towards evading flg22 variants: (1) Enhancing FLS2-flg22 interaction around key evasion sites to recognize weak agonistic flg22 variants with reduced binding affinity to FLS2. (2) Strengthening direct FLS2-BAK1 interaction to enable recognition of antagonistic flg22 variants that hinder BAK1 recruitment. (3) Harnessing ‘dark matter’ polymorphisms that enhance performance through mechanisms that are not yet fully understood. **d,** Workflow for FLS2 engineering using these design principles. Based on the type of evasion—antagonism or loss of interaction with FLS2—appropriate design principles can be applied. Data are presented as mean ± SEM. The working concentration of elicitors is 100 nM.

Transplantation of LRR20-EJM^GmFLS2b^ enabled SlFLS2 and NbFLS2 to recognize flg22^Rso^, but not AtFLS2, VvFLS2, and VrFLS2XL (**Fig. 5b**). To explain this, we assessed the evasion and antagonistic effects of flg22^Rso^ across different FLS2 orthologs (**Extended Data Fig. 5**). Flg22^Rso^ acted as a strong antagonist for SlFLS2 and NbFLS2, but showed minimal antagonistic effects for AtFLS2, VvFLS2, and VrFLS2XL. The correlation between the antagonism by flg22^Rso^ and the ability to gain flg22^Rso^ recognition after LRR20-EJM^GmFLS2b^ transplantation suggests that strengthening direct FLS2-BAK1 interaction is particularly effective for flg22 variants that function as strong antagonists. This finding also underscores the importance of first determining the evasion mechanism before initiating receptor engineering (**Fig. 5c, d**).

## Discussion

During plant-pathogen coevolution, plants have evolved PRRs with novel recognition specificities to detect PAMPs of pathogens, and some pathogens have evolved polymorphic PAMPs to evade the recognition by PRRs. Editing native PRRs to gain recognition towards evading PAMPs offers a feasible strategy to generate durable broad-spectrum disease resistance without causing legal issues linked to transgene-based PRR transfer, but our knowledge on how to engineer such PRRs remains limited. In this study, we reverse-engineered two natural FLS2 variants and discovered fundamental design principles of broader recognition spectra against evading flg22 epitopes.

In our first reverse-engineering case using VrFLS2XL, we discovered that flg22 variants with reduced binding affinity to FLS2 can be addressed by enhancing the FLS2-flg22 interaction around key evasion sites of flg22. While this strategy seems straightforward and is supported by several studies^20, 25^, it might not be easy to implement. Our results showed that approximately ten residue changes are required to make VvFLS2 respond to flg22^Atum^ (**Fig. 2f**). Some of these residues are predicted to directly interact with key evasion sites of flg22^Atum^, some seem to interact with flg22^Atum^ residues adjacent to key evasion sites to assist with ligand binding, and others, including buried residues, appear to stabilize local conformation (**Fig. 2b-c**). Notably, buried residues also play crucial roles, which is overlooked in Repeat Conservation Mapping (RCM) analysis^45^ and previous attempts to engineer FLS2^20^. Moreover, the occasional simultaneous decreases in flg22^Pa^ and flg22^Atum^ responses observed in our mutation analysis (**Fig. 2d**, **Extended Data Fig. 2c**) suggests that the novel recognition specificity might come at the expense of existing recognition capabilities, although this could be rescued. Overall, the novel flg22^Atum^ recognition capability of VrFLS2XL appears to involve a complex and finely tuned interaction network. Therefore, further research is needed to elaborate this design principle to develop simpler solutions. Additionally, while we derived this engineering strategy by studying the weakly agonistic flg22^Atum^, its applicability for engineering recognition of antagonistic flg22 variants remains to be investigated.

The reverse engineering of GmFLS2b offers a strategy to counteract the antagonistic flg22^Rso^ by strengthening the direct FLS2-BAK1 interaction without altering any flg22-interacting residues. Notably, the gain of flg22^Rso^ recognition by GmFLS2a was achieved with the modification of only two residues (**Fig. 4c-h**), suggesting the effectiveness of this strategy. However, this strategy works only for FLS2 homologs to which flg22^Rso^ acts as an antagonist (**Fig. 5b** and **Extended Data Fig. 5**). Although this strategy was not reported in the initial GmFLS2b work^25^, the importance of FLS2-BAK1 direct interaction was suggested by several previous studies. For example, transplanting LRR19-24 of SlFLS2 to AtFLS2 enabled AtFLS2 to respond to flg22^Pa^ with 18AYA20 mutations^33^, and transplanting LRR18-24 of SlFLS2 to AtFLS2 led to AtBAK1-dependent autoactivation^46^. Future research should explore whether this principle applies to engineering recognition of antagonistic flg22 variants beyond flg22^Rso^. These findings also imply that the flg22-mediated and the direct FLS2-BAK1 interaction interfaces function coordinately rather than sequentially to recruit BAK1 since the reduced interaction at the first interface can be compensated by an increased interaction at the second interface.

Using DNA shuffling, we discovered ‘dark matter’ polymorphisms outside the flg22- or BAK1-interacting interfaces of GmFLS2a/b that significantly affect flg22^Rso^ recognition through unknown mechanisms, without altering FLS2 accumulation (**Fig. 3c-e** and **Fig. 4b, i-l**). The three asparagine-related polymorphisms within LRR12-19^GmFLS2b^ might lead to gains and losses in glycosylation. Although FLS2 has been shown to tolerate changes in glycosylation status^47, 48, 49^, it is possible that glycosylation subtly modulates FLS2’s recognition specificity through various mechanisms^50^. The six ‘enhancer’ polymorphisms located in the C-terminal LRR capping domain and extracellular juxtamembrane domain point to the significance of this previously overlooked region, which aligns with earlier findings that mutations of cysteines within this region enhance FLS2-mediated ROS production^47^. ‘Dark matter’ polymorphisms may not only influence FLS2 itself but could also affect its interaction with other positive or negative regulators, which could be further explored by immunoprecipitation-mass spectrometry assays in the future. Further investigation on ‘dark matter’ polymorphisms will deepen our understanding of PRRs and may reveal general, ligand-independent design principles to engineer PRRs with enhanced functionality.

From a methodological standpoint, our study establishes a pipeline for the identification of minimal sequence determinants responsible for novel recognition specificities in natural PRR variants. The three approaches used – domain swapping, computational design, and DNA shuffling – each have their advantages and limitations, but they complement each other. Domain swapping is well-suited for rough mapping of key residues. While finer mapping is possible by making more precise segmentations, this significantly increases the workload for molecular cloning. Computational design works best in cases where structures are resolved or predicted with high confidence. The more we understand about the PRR-ligand pair, the easier the mapping process becomes. However, this method may overlook crucial polymorphisms that are beyond current knowledge, as seen in the case of GmFLS2a/b. DNA shuffling, on the other hand, enables rapid, unbiased fine mapping without much prior knowledge. However, it requires parental DNA fragments with high homology and polymorphic sites that are not tightly clustered^51^. For example, LRR20-EJM fragments of GmFLS2a and GmFLS2b are suitable candidates to shuffle, whereas LRR12-19 fragments of VvFLS2 and VrFLS2XL are not due to the large number of densely clustered polymorphic sites. Beyond these three approaches, other strategies can be employed, such as leveraging coevolutionary insights^52^, ancestral sequence reconstruction^53^, and using Golden Gate Shuffling that combines the advantages of domain swapping and DNA shuffling^54^. Additionally, our study introduces an optimized method for constructing large, high-quality libraries of DNA variants directly in *A. tumefaciens*, using Golden Gate cloning and the *SacB* counterselection marker (**Fig. 4a**). However, the relatively low screening throughput remains a bottleneck, calling for technological innovation.

In addition to providing design principles for future FLS2 improvement, the engineering outcomes of this study already hold potential for practical applications. For example, to generate crown gall-resistant grapes, introducing 14 residue changes from the ‘comp.Med’ list (**Fig. 2a**) into VvFLS2 requires 18 nucleotide substitutions, which is below the threshold of 20 nucleotides set by the current proposed EU regulation on plants produced by certain new genomic techniques (COM/2023/411). These modifications could be introduced by prime editing and base editing, both of which are already available for grapevine^55, 56^, or by homologous recombination^57^, which seems more straightforward but has not been reported for grapevine yet. Given that wild-type GmFLS2b has a relatively low recognition capacity for flg22^Rso^ compared to flg22^Pa^, yet still confers decent resistance against *R. solanacearum* when transiently expressed in roots under the control of the strong constitutive 35S promoter^25^, we assume that our ‘superGmFLS2’ engineered variant could confer an even stronger resistance when stably transformed into crops with an appropriate promoter. Alternatively, it could serve as a template for engineering native FLS2s in other crops to generate resistance to bacterial wilt disease.

Together, our study establishes fundamental principles for engineering expanded recognition spectra against two classic types of evading flg22 variants, provides a methodology for identifying sequence determinants of novel recognition specificity from natural PRR variants, and offers ready-to-use genetic resources for developing resistance to crown gall disease and bacterial wilt. Combined with advancements in protein engineering techniques, genome editing technologies, and evolving legal regulations as well as public acceptance towards genome-edited crops, our work paves the way for utilizing PRR engineering to create gene-edited crops with durable broad-spectrum disease resistance.

## Methods

### Plant materials and synthetic peptides

Five-week-old *Nicotiana benthamiana fls2* mutant^26^ was used for ROS burst assays and infection assays with *A. tumefaciens* carrying GUS. Transgenic *N. benthamiana* expressing aequorin^58^ was used for cytoplasmic calcium measurement. *N. benthamiana* plants were grown in the greenhouse at 60 % relative humidity and 25 °C/22 °C in a 16-hour/8-hour day/night cycle. Synthetic peptides of flg22^Pa^, flg22^Atum^, and flg22^Rso^ (sequences shown in main figures) were produced by Scilight Biotechnology (Beijing, China) at >80 % purity. Peptide powder was dissolved in sterile ddH_2_O to a concentration of 10 mM as a stock solution and stored at -20 °C. Working solution with a certain concentration was freshly diluted from the stock solution before use.

### Transient expression in *N. benthamiana*

*A. tumefaciens* GV3101 transformed with certain binary plasmid was grown overnight in liquid LB medium supplemented with gentamycin, rifampicin, and antibiotics corresponding to the binary vector (spectinomycin for FLS2 plasmids, kanamycin for P19 and GUS plasmids). Cells were collected by centrifugation (3000 RCF, 5min), washed once, and resuspended in the infiltration buffer (10 mM MES-KOH, pH 5.8, 10 mM MgCl_2_, 150 μM acetosyringone). Cell density (OD_600_) was measured using a Bio-Rad SmartSpec™ Plus spectrophotometer. For infiltration, *A. tumefaciens* carrying the FLS2 plasmid and a second strain carrying the P19 plasmid were mixed at a 4:1 ratio to reach a final OD_600_ of 0.5 (0.4 FLS2 + 0.1 P19). For a quantitative comparison of two FLS2 variants, two freshly transformed *A. tumefaciens* strains carrying two constructs were symmetrically infiltrated into two sides of the midrib.

### Molecular cloning

To generate FLS2 chimeras and mutants in a modular manner, a unified multi-kingdom Golden Gate cloning platform was used^36, 59^. Template plasmids of NbFLS2, AtFLS2, and VvFLS2 were from the lab stock. VrFLS2XL and SlFLS2 were gifts from Georg Felix (University of Tübingen). GmFLS2a and GmFLS2b were gifts from Alberto Macho (Chinese Academy of Sciences). N-LRR11, LRR12-19, LRR20-EJM, and TMD+ICD segments of aforementioned FLS2s were amplified from template plasmids and subcloned to p641-Esp3I/BpiI level 1 entry vectors. To assemble FLS2 segments, a level 2 receiver plasmid was constructed by assembling a 35S promoter (Addgene #54406), a *LacZ* expression cassette flanked by Esp3I sites (Addgene #54458), a C-terminal mEGFP tag (Addgene #54410 with A206K mutation), a NOS terminator (Addgene #54407), and dummies (Addgene #54342, #54345) into a level 2 binary vector (Addgene #54346). LRR12-19^VvFLS2^ fragments carrying computationally predicted polymorphic residue sets (‘comp.Max’, ‘comp.Med’, ‘comp.Min’), the LRR20-EJM^GmFLS2a^ fragment carrying YHR and YHAIR (F579Y, Q600H, S603A, M626I, H723R) mutations, and the codon-optimized LRR20-EJM^GmFLS2b^ fragment for DNA shuffling were synthesized by Twist Bioscience. To construct the receiver plasmid for shuffled DNA fragments, the *SacB* expression cassette was first cloned into the p641-BpiI level 1 vector using pT18mobsacB (Addgene #72648) as PCR template, meanwhile introducing flanking Esp3I sites and the same overhangs as the LRR20-EJM fragment. Then, the level 2 receiver plasmid was constructed by assembling level 1 plasmids of 35S promoter, NOS terminator, C-terminal mEGFP tag, N-LRR11^GmFLS2a^, LRR12-19^GmFLS2a^, TMD+ICD^GmFLS2a^, *SacB* expression cassette and dummies. Site-directed mutagenesis was conducted in level 1 plasmid using a Golden Gate-based strategy^60^. DNA sequences were listed in **Supplementary Table 1**. Primer sequences were listed in **Supplementary Table 2**.

### Measurement of ROS production

At least eight leaf discs per treatment were harvested from each leaf area transiently expressing a certain FLS2 variant were harvested with a Ø 4 mm biopsy punch (KAI, #BP-40F) and floated on 100 μL distilled water overnight in white 96-well plates (Greiner Bio-One, #655075) in darkness. Immediately after replacing water with ROS assay solution (100 μM luminol (Sigma, #A8511), 20 μg/mL horseradish peroxidase (Sigma, #77332)) with or without elicitors, luminescence was recorded using a microplate reader (Tecan Spark® or Tecan Infinite® Lumi) with 1-min intervals and 250-ms integration time. At least three different leaves from three plants were sampled for each FLS2 variant.

### Cytoplasmic calcium measurement

At least eight leaf discs per treatment were harvested from each leaf area transiently expressing a certain FLS2 variant using a Ø 4 mm biopsy punch (KAI, #BP-40F) and incubated in 100 μL of 10 μM coelenterazine water solution in darkness overnight in white 96-well plates (Greiner Bio-One, #655075). After replacing the coelenterazine solution with 75 μL of distilled water, luminescence was measured using a Tecan Spark® multimode microplate reader. As a baseline, luminescence was first recorded in 30-second intervals for 5 minutes. Then, 25 μL of water (as Mock) or a four-time concentrated elicitor water solution was added to each well, and luminescence was recorded in 30-s intervals for 30 min. The remaining AEQ signal was discharged by adding 100 μL of discharge solution (2 M CaCl_2_, 20 % EtOH) and measured for 90 s. A normalized signal ‘L/L_max_’ was calculated for each time point, where ‘L’ is the recorded absolute luminescence at each time point and ‘L_max_’ is the total luminescence during 90 s discharging.

### Protein extraction and western blot

For total protein extraction, *N. benthamiana* leaf tissues were sampled into 2 mL tubes with Ø 4 mm glass beads, snap-frozen in liquid nitrogen, grounded in SPEX SamplePrep 2010 Geno/Grinder®, and homogenized in protein extraction buffer (50 mM Tris-HCl pH 7.5, 100 mM NaCl, 10 % glycerol, 2 mM EDTA, 5 mM DTT, 1 % IGEPAL CA630, 2 mM Sodium molybdate, 1 mM Sodium fluoride, 1 mM Sodium orthovanadate, 4 mM Sodium tartrate, protease inhibitor cocktail). Samples were incubated at 4 °C for 10-20 min and then centrifuged at 16,000 g for 20 min at 4 °C to remove debris. Supernatants were mixed with SDS loading buffer and boiled at 95℃ for 5 minutes before loading. Proteins were separated using 10% SDS-PAGE gels and transferred to a PVDF membrane. The membrane was blocked using 5 % skim milk in TBST and probed using the corresponding antibodies diluted in TBST-5 % skim milk. Blotted membranes were stained with Coomassie brilliant blue (CBB) as loading control. To detect the expression levels of FLS2 variants, GFP Antibody (B-2) (Santa Cruz, #sc-9996) was used as the primary antibody, and Anti-Mouse IgG (Fc specific)–Peroxidase (Sigma-Aldrich, #A0168) was used as the secondary antibody.

### Detection of MAPK activation

After transiently expressing FLS2 variants for 3 days, intact *N. benthamiana* leaves were syringe-infiltrated with 100 μM flg22^Rso^ or ddH_2_O only as a mock control. After flg22^Rso^ treatment for 15 min, eight leaf discs were harvested into 2-mL tubes containing five Ø 4-mm glass beads. using a Ø 8 mm biopsy punch (KAI, #BP-80F). Protein extraction, SDS-PAGE, and western blot were performed as aforementioned, but 5% BSA (bovine serum albumin) in TBST was used for blocking and diluting antibodies. Phospho-p44/42 MAPK (Erk1/2) (Thr202/Tyr204) antibody (Cell Signaling Technology, #9101) was used to detect phosphorylated MAPKs.

### Measurement of GUS activity by histochemical and fluorometric assays

After transiently expressing FLS2 variants for 3 days, whole *N. benthamiana* leave was infiltrated with *Agrobacterium tumefaciens* GV3101 carrying pBIN19g:GUS (intronic) plasmid at OD = 0.5 (suspended in infiltration buffer without acetosyringone). For the histochemical GUS assay, 4 days after infiltration, the whole leave was collected and vacuum-infiltrated thoroughly in GUS staining solution (0.1 M sodium phosphate buffer pH 7.0, 10 mM EDTA, 0.1 % Triton X-100, 1 mM K_3_[Fe(CN)_6_], 2 mM X-Gluc). After overnight incubation in GUS staining solution at 37 °C, samples were destained in 70 % ethanol at 60 °C before taking photos. For the fluorometric GUS assay, 4 days after infiltration, six leaf discs for each FLS2 variant were sampled in 2-mL tubes containing three Ø 4-mm glass beads using a Ø 4-mm biopsy punch (KAI, #BP-40F) and were snap-frozen in liquid nitrogen. Subsequent steps were performed as previously described^61^.

### Prediction, visualization, and analysis of protein complex structure

The structure of the FLS2-flg22-BAK1 complex was predicted using AlphaFold3 server^62^. The top-ranked (by ipTM values) models were further analyzed using Mapiya^63^ to determine all possible interactions, e.g., hydrogen bonds, salt bridges, hydrophobic contacts, electrostatic interactions (dipole-dipole, dipole-π stacking), within 4 and 8 angstroms between FLS2 and flg22 or FLS2 and BAK1. The predicted models and interactive residues were visualized by PyMol^64^.

### DNA shuffling and direct variant library construction in *A. tumefaciens*

For details and tips to plan a DNA shuffling experiment, please refer to Meyer et al.^51^. Parental DNA fragments for DNA shuffling were prepared by amplifying level 1 entry plasmids carrying LRR20-EJM^GmFLS2a^ and LRR20-EJM^GmFLS2b^ using the outer primer pair SZ172 and SZ173, and purifying PCR product from the 1% agarose gels. For an optimal DNA shuffling outcome, LRR20-EJM^GmFLS2b^ was codon-optimized to achieve maximum homology with LRR20-EJM^GmFLS2a^. DNA Shuffling was performed using the JBS DNA-Shuffling Kit (Jena Bioscience, #PP-103). Shuffled DNA fragments were amplified using the inner primer pair SZ621 and SZ622. The PCR product was purified and ligated into the *SacB*-containing vector by Golden Gate cloning. Golden Gate product was purified using the DNA Clean & Concentrator-5 kit (Zymo Research, #D4003) and transformed to *A. tumefaciens* GV3101 by electroporation. Transformed cells were plated on LB plates supplemented with gentamycin, rifampicin, spectinomycin, and 5% sucrose. Primer sequences were listed in **Supplementary Table 2**.

### Analysis of DNA shuffling results

The performance of a shuffled variant is defined by the ratio of the peak values (RLU_max_) of flg22^Rso^- and flg22^Pa^-induced ROS curves. The performance of a residue at a polymorphic site is calculated as the average performance of all variants carrying this residue. For example, the performance of alanine from GmFLS2a at position 536 is calculated by averaging the performances of all shuffled variants with an alanine at position 536. A Welch’s t-test (unequal variances t-test) is conducted for each polymorphic site to compare the mean performance of residue from GmFLS2b against that from GmFLS2a. A high T-statistic absolute value suggests a significant difference between the performances of residues sourced from GmFLS2a and GmFLS2b at that position, implying that this site might be important for flg22^Rso^ recognition. Instead, a low T-statistic absolute value suggests that this site might not be important for flg22^Rso^ recognition. A positive T-statistic value means that the residue of GmFLS2b origin at this position positively contributes to flg22^Rso^ recognition. A negative T-statistic value means that the residue of GmFLS2a origin at this position negatively contributes to flg22^Rso^ recognition.

## Acknowledgments

We thank all members of the Zipfel group for discussions – in particular Kyle Bender, Oliver Johanndrees, Gijeong Kim, Moutassem Omary, Simon Snoeck, and Keran Zhai – as well as Julia Santiago, Gabriel Scalliet and Julia Vorholt. We thank Qiang Cheng for the kind gift of *N. benthamiana fls2* mutant seeds, Georg Felix for plasmids containing VrFLS2XL and SlFLS2, and Alberto Macho for plasmids containing GmFLS2a and GmFLS2b. We thank UZH-BIO286 students Charles Guillain and Julian Kamber for helping with the experiments. This work was funded by the University of Zurich (C.Z. and A.C.) and a Zurich-Basel Plant Science Center-Syngenta fellowship (to S.Z. and C.Z.).

## Author contributions

S.Z. and C.Z. conceived the project and designed the experiments. C.Z. and A.C. supervised the project and acquired funding. S.Z. and H.L. performed experiments and data analysis. S.L. performed structural modeling and analyzed the DNA shuffling results. S.Z. and S.L. made figures. S.Z. and C.Z. wrote the manuscript with inputs from all the authors.

## Extended Data Information

**Extended Data Fig. 1:**
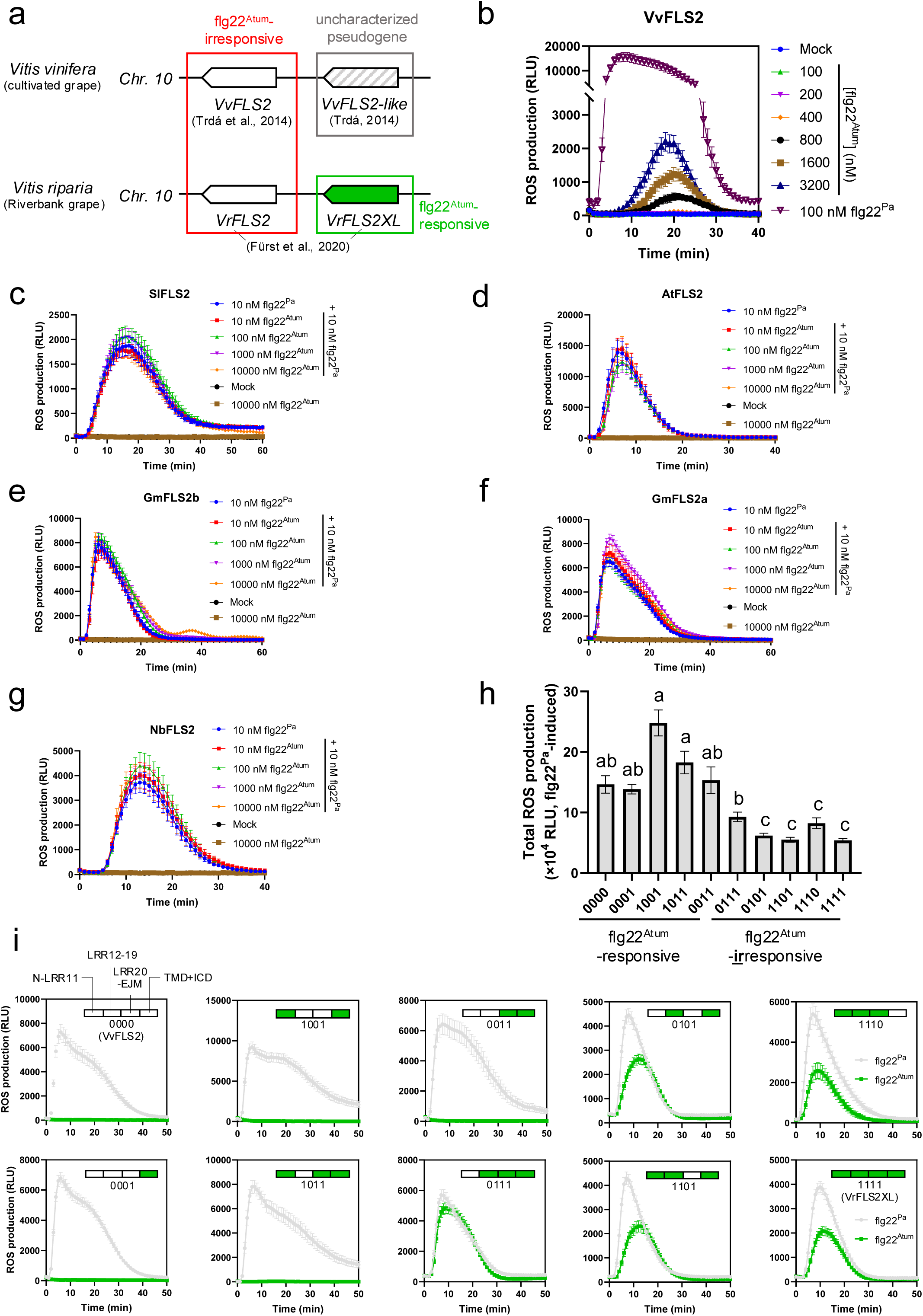
Evasion of recognition by flg22^Atum^ across different FLS2 homologs and flg22^Atum^ responsiveness of VvFLS2-VrFLS2XL chimeras (related to Fig. 1). **a,** Summary of the reported flg22^Atum^ responsiveness of FLS2 homologs in *Vitis vinifera* and *Vitis riparia*. Chr.: chromosome. **b,** Kinetics of VvFLS2’s ROS responses to flg22^Pa^ and increasing concentrations of flg22^Atum^ (related to Fig. 1b). **c-g,** Characterization of flg22^Atum^ evasion and its potential antagonistic effect on SlFLS2 (**c**), AtFLS2 (**d**), GmFLS2a (**e**), GmFLS2b (**f**), NbFLS2 (**g**). No significant differences in total ROS production were observed between treatments in panels **c–f**. **h,** Total flg22^Pa^-induced ROS production of chimeric receptors. **i,** ROS burst kinetics of chimeric receptors in response to flg22^Pa^ and flg22^Atum^. Unless specified, 100 nM of each elicitor was used to trigger responses, with total ROS measured as relative light units (RLU) accumulated over 40 minutes. Data are shown as mean ± SEM. Kruskal-Wallis test followed by Dunn’s multiple comparisons test was used for statistical analysis in panel **h**. Significant differences at *P* < 0.05 are indicated by letters.

**Extended Data Fig. 2:**
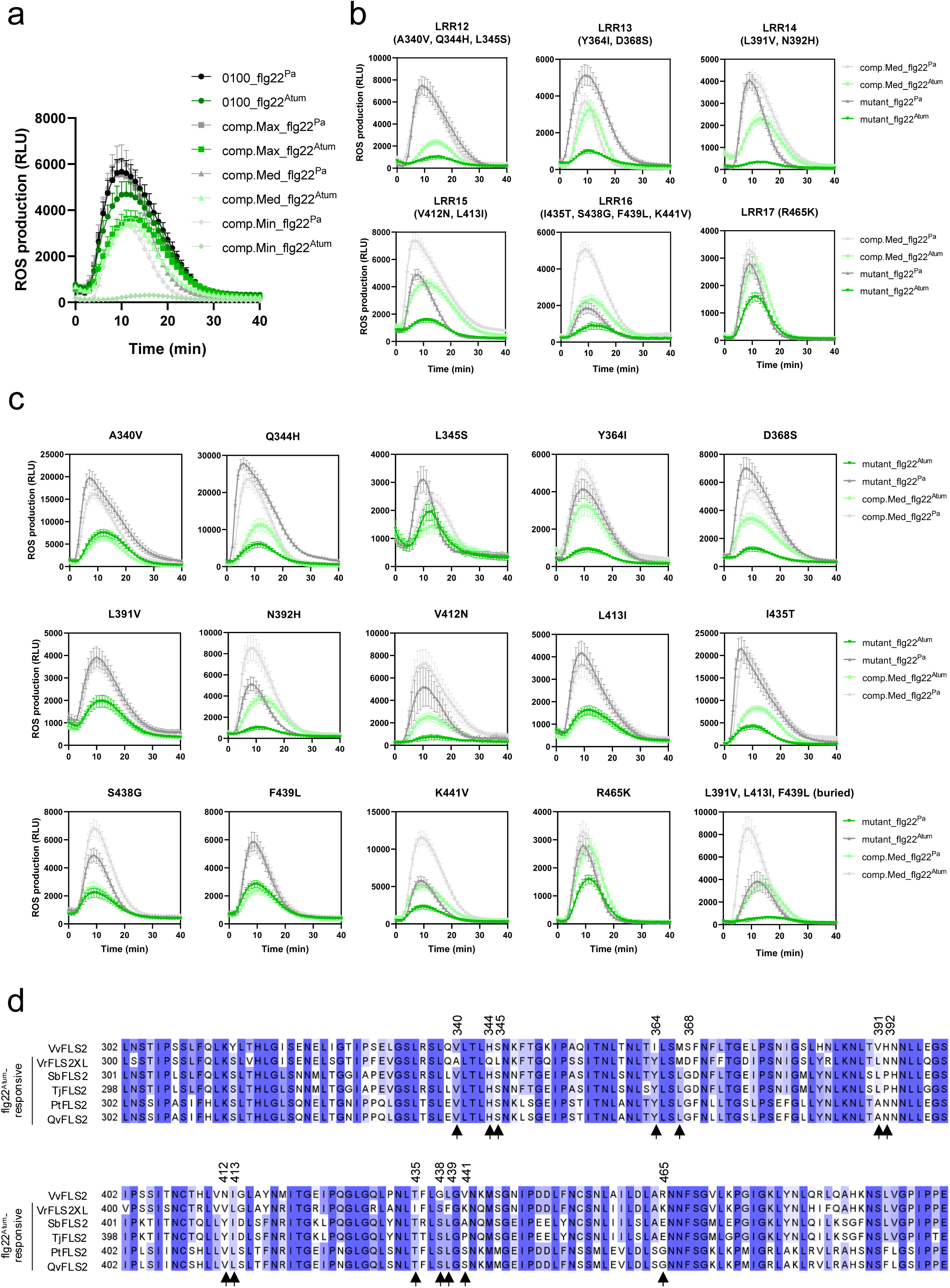
Characterization of computationally designed variant ‘comp.Med’ (related to Fig. 2). **a,** Flg22^Pa^- and flg22^Atum^-induced ROS production of engineered FLS2 variants listed in Fig. 2a (corresponding to Fig. 2d). **b,** Effects of mutating clusters of polymorphic residues in each LRR from LRR12-17 of ‘comp.Med’. **c,** ROS burst kinetics of ‘comp.Med’ with single or multiple mutations at polymorphic sites (corresponding to Fig. 2f). **d,** Sequence alignment of the LRR12-19 region of reported flg22^Atum^-responsive FLS2 variants. Polymorphic sites within ‘comp.Med’ are indicated with arrows and their positions within VrFLS2XL. (Qv, *Quercus variabilis*; Tj, *Trachelospermum jasminoides*; Sb, *Salix babylonica*; Pt, *Populus trichocarpa*). Data are presented as mean ± SEM. Sequence alignment and visualization were performed using Jalview^65^. Refer to Fig. 2 for further details.

**Extended Data Fig. 3:**
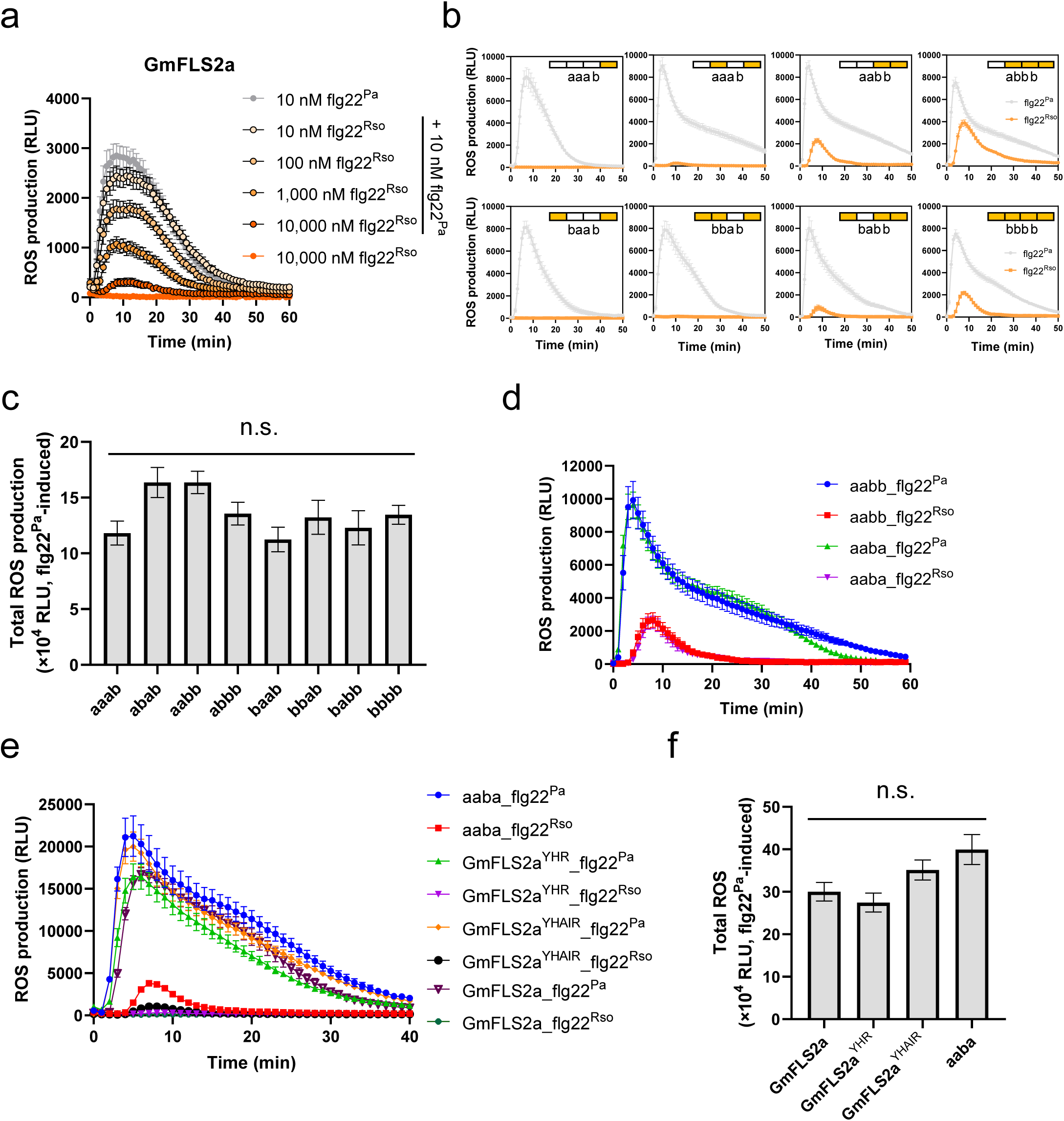
ROS burst kinetics and total flg22^Pa^-induced ROS production of GmFLS2a/b chimeras and GmFLS2a variants with predicted gain-of-recognition polymorphic residues (related to Fig. 3). **a,** ROS burst kinetics corresponding to Fig. 3b. **b-c,** ROS burst kinetics (**b**) and total flg22^Pa^-induced ROS production (**c**) of GmFLS2a/b chimeras (related to Fig. 3c). **d,** Flg22^Pa^- and flg22^Rso^-induced ROS responses of ‘aabb’ and ‘aaba’ chimeras. **e-f,** ROS production kinetics (**e**), and total flg22^Pa^-induced ROS production (**f**) of GmFLS2a variants carrying predicted gain-of-recognition residues (related to Fig. 3g). Data are presented as mean ± SEM. Kruskal-Wallis test followed by Dunn’s multiple comparisons test was used for statistical analysis of **c** and **f**. ‘n.s.’ indicates no significant difference. Refer to Fig. 3 for further details.

**Extended Data Fig. 4:**
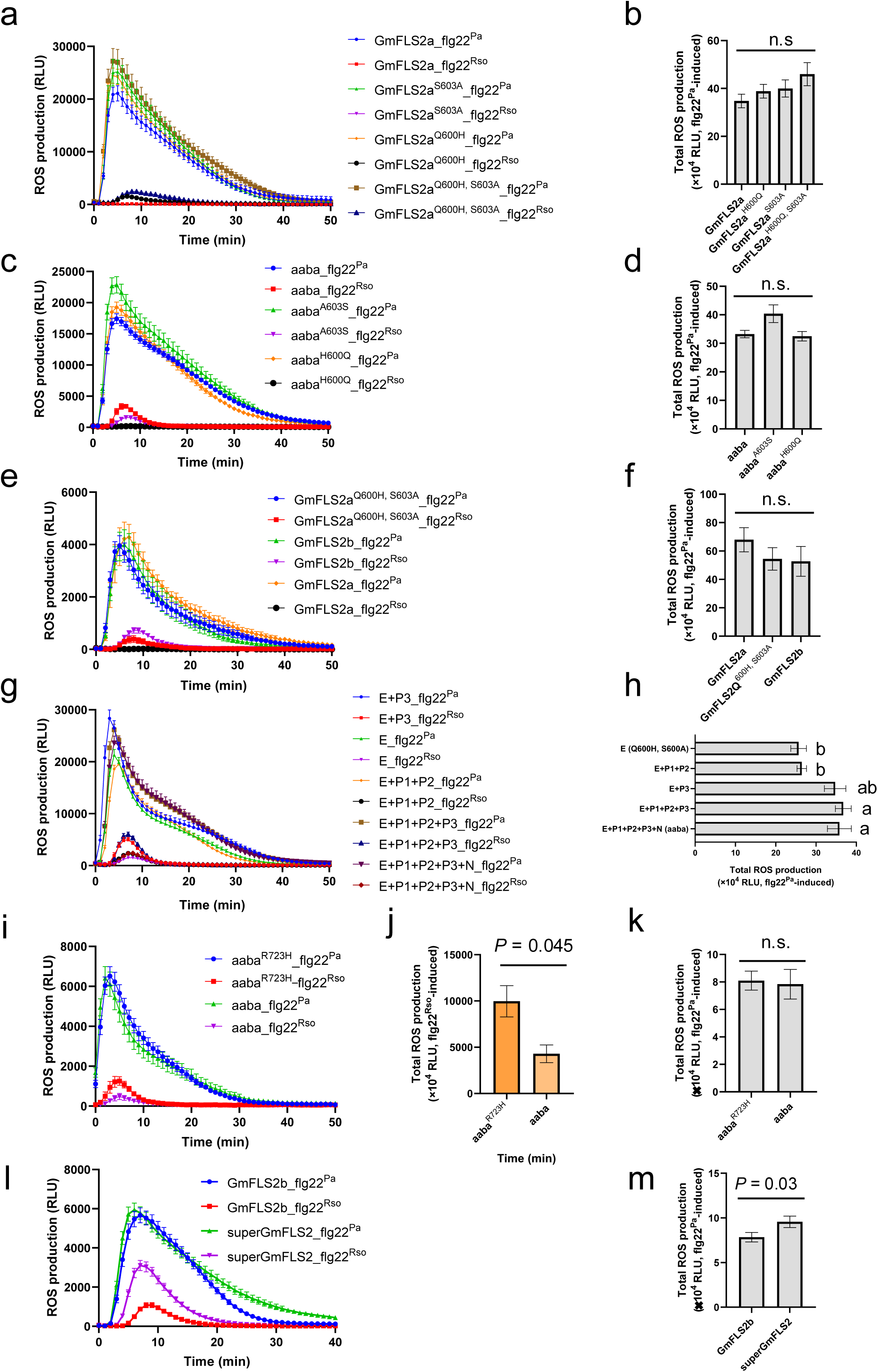
Characterization of minimal gain-of-function residues, along with ‘enhancer’ and ‘suppressor’ polymorphisms predicted by DNA shuffling (related to Fig. 4). **a-b,** ROS production kinetics (**a**) and total flg22^Pa^-induced ROS production (**b**) of GmFLS2 variants characterized in Fig. 4c. **c-d,** ROS production kinetics (**c**) and total flg22^Pa^-induced ROS production (**d**) of GmFLS2 variants characterized in Fig. 4d. **e-f,** ROS burst kinetics (**e**) and total flg22^Pa^-induced ROS production (**f**) of GmFLS2 variants characterized in Fig. 4e. **g-h,** ROS burst kinetics (**g**) and total flg22^Pa^-induced ROS production (**h**) of GmFLS2 variants characterized in Fig. 4i. **i-k,** Effects of R723H on flg22^Pa^- and flg22^Rso^-induced ROS production. **l-m,** ROS burst kinetics (**l**) and total flg22^Pa^-induced ROS production (**m**) of GmFLS2 variants characterized in Fig. 4j. Data are presented as mean ± SEM. Kruskal-Wallis test followed by Dunn’s multiple comparisons test was performed for **b**, **d**, **f**, and **h**, and significant differences at *P* < 0.05 are indicated by letters. Mann-Whitney test was performed for **j**, **k**, and **m**. The ‘n.s.’ denotes no significant difference. For additional details, refer to Fig. 4.

**Extended Data Fig. 5:**
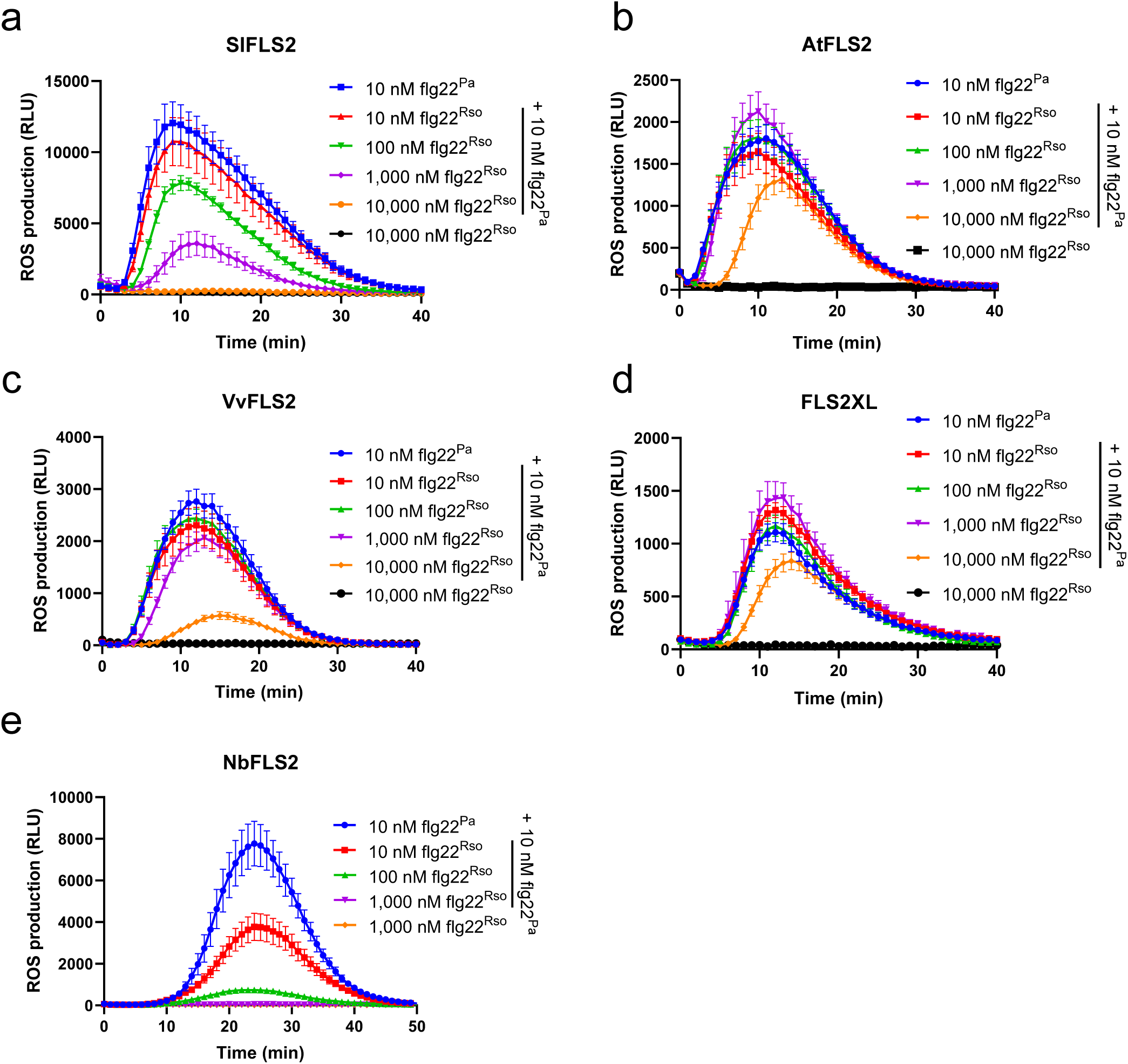
Evasion and antagonistic effect of flg22^Rso^ across FLS2 homologs (related to Fig. 5). **a,** SlFLS2. **b,** AtFLS2. **c,** VvFLS2. **d,** VrFLS2XL. **e,** NbFLS2. Data are presented as mean ± SEM. The working concentration of elicitors is 100 nM.

**Supplementary Table 1. DNA sequences of natural FLS2 fragments and synthesized FLS2 fragments. BsaI cutting sites are underlined.**

**Supplementary Table 2. Primers used in this study.**

**Supplementary Table 3. Shuffled GmFLS2a/b LRR20-EJM variants used for analysis.**

## Notes

### Competing Interest Statement

The authors have declared no competing interest.

